# Age-related decline in behavioral discrimination of amplitude modulation frequencies compared to envelope-following responses

**DOI:** 10.1101/193268

**Authors:** Jesyin Lai, Edward L. Bartlett

## Abstract

The ability to discriminate modulation frequencies is important for speech intelligibility because speech has amplitude and frequency modulations. Neurophysiological responses assessed by envelope following responses (EFRs) significantly decline at faster amplitude modulation frequencies (AMF) in older subjects. A typical assumption is that a decline in EFRs will necessarily result in corresponding perceptual deficits. To test this assumption, we investigated young and aged Fischer-344 rats’ behavioral AMF discrimination abilities and compared to their EFRs. A modified version of prepulse inhibition (PPI) of acoustic startle reflex (ASR) was used to obtain behavioral performance. A PPI trial contains pulses of sinusoidal AM (SAM) at 128 Hz presented sequentially, a SAM prepulse with different AMF and a startle-eliciting-stimulus. To account for hearing threshold shift or age-related synaptopathy, stimulus levels were presented at 10-dB lower or match to the aged peripheral neural activation (using auditory brainstem response wave I amplitude). When AMF differences and modulation depths were large, young and aged animals’ behavioral performances were comparable. Aged animals’ AMF discrimination abilities declined as the AMF difference or the modulation depth reduced, even compared to the young with peripheral matching. Young animals showed smaller relative decreases in EFRs with reduced modulation depths. The correlation of EFRs and AM perception was identified to be more consistent in young animals. The overall results revealed larger age-related deficits in behavioral perception compared to EFRs, suggesting additional factors that affect perception despite smaller degradation in neural responses. Hence, behavioral and physiological measurements are critical in unveiling a more complete picture on the auditory function.

## 1. Introduction

Presbycusis is common and unavoidable in the elderly due to its properties of chronic deterioration and is asymptomatic early in life [66, 20]. It has been reported as the third most prevalent chronic disorder in the elderly (*≤* 65 years old) after hypertension and arthritis in the United States [40]. Age-related changes in auditory structures and functions exist in both the peripheral and central auditory systems [58, 59, 66, 6, 18, 72]. Age-related degradation of the auditory periphery comprises loss or dysfunction of the inner and outer hair cells [24, 59], alterations in the stria vascularis leading to endocochlear potential reduction [8], and/or diminished auditory nerve fibers (ANFs) and synapses [60]. Meanwhile, changes in excitatory/inhibitory balance are reported and described as one of the main causes of age-related auditory deficits in the central auditory system [6, 7, 53, 46]. Auditory central degradation could result in degraded processing of complex sounds especially in challenging situations, for example speech recognition in a cocktail party [22].

Human speech consists of complex and rapid modulations in amplitude and frequency over time that are crucial for precise speech recognition [54, 61, 75]. Previously, our research team and others have revealed significant age-related differences in temporal processing, assessed physiologically by envelope following responses (EFRs) at the levels of the auditory midbrain and brainstem, at faster AM frequencies (AMFs) [47, 52]. Psychoacoustic studies using temporal modulation transfer functions (tMTFs) have also shown that older adults have poor periodicity coding due to higher thresholds in modulation depth and frequency modulation (FM) detection [25, 26]. We have collected neurophysiological evidence from young and aged rats showing age-related differences in temporal processing of AM and FM [48, 47]. It is assumed that larger EFR responses elicited by AM sounds are associated with better perceptual performance [48, 2, 43]. However, there is a lack of behavioral evidence that clarifies and confirms the relationship of physiological and behavioral responses.

To assess and determine changes in neural processing related to auditory impairments or brain disorders, the acoustic startle response (ASR) with its modulation by a non-startling prepulse is broadly applied in behavioral sensory studies [37, 14, 62]. The ASR is a type of reflexive behavior manifested as a transient contraction of facial and skeletal muscles in respond to a sud-den, brief and intensely loud sound [64, 39]. In rats, the ASR can be elicited by an acoustic stimulus that is approximately more than 80 dB above the hearing threshold [50]. Therefore, measurement of ASR can be used as an indicator for the behavioral responsiveness or perception to acoustic stimuli. Startle reflex behavior is convenient for age-related auditory studies because it is an unconditioned reflex reaction and no animal training is required. It has also been demonstrated that the ASR can be measured at any age past juvenile in rats [67, 69]. The primary ASR circuit comprises the cochlear root neurons, neurons in the caudal pontine reticular nucleus (PnC) and spinal motor neurons [36, 10, 21]. This simple neural circuit has extremely short latency because it involves only a few synapses located in the lower brainstem [36, 10].

The amplitude and probability of a startle movement following a SES can be modulated by non-startling prepulses. A prepulse is a stimulus presented prior to the SES. The amplitude of the ASR is attenuated significantly when the prepulse is detected and processed by the subject [13]. Therefore, inhi-bition of the startle reaction using a prepulse is termed prepulse inhibition (PPI). The magnitude of PPI is proportional to the subject’s detectability of the prepulse [33]. Prepulses have been used in the forms of acoustic [29], visual [4] and tactile [51]. Animal studies have shown that auditory PPI is associated with the function of the cochlear nucleus, the inferior and superior colliculi (I/SC) and the pedunculopontine tegmental nucleus [36]. When a prepulse is presented, the signal travels from the level of the cochlea to the IC and then travels collaterally to the SC. Subsequently, the SC excites the pedunculopontine tegmental nucleus, which inhibits the PnC, resulting in reduced startle response [13, 36]. Hence, an interval of 20-500 ms between the prepulse and the SES should provide sufficient time for the signal to inhibit the ASR via PnC inhibition [13, 36, 37].

PPI can be induced by prepulses with various temporal characteristics. Prepulse duration up to 100 ms are generally used in most PPI experiments [32, 31, 17, 65]. Recently, other applications of the PPI paradigm were de-veloped using complex modulatory stimuli with relatively long duration, for example 50-1000 ms gap prepulses in background noise [62]. Detection of an amplitude modulated prepulse, which was presented during 1 s before the SES, from a background of unmodulated noise has been demonstrated in gerbils of two-month age [41]. Speech sounds of 100-300 ms have also been used as prepulses in rats [15, 16]. Floody and Kilgard (2007) showed that Sprague-Dawley rats of approximately four-month age were able to distinguish syllable [pae] from [bae] with the application of the PPI paradigm.

In this study, we investigated AMF discrimination abilities of young and aged F344 rats using the PPI paradigm. A modified test paradigm, adapted from Floody and Kilgard’s (2007) speech discrimination tasks, was applied by replacing speech sounds with AM sounds. AM sounds modulated with AMFs different from the AMF of background sounds were used as prepulses. The behavioral results were then compared to EFRs of tMTFs recorded from each of the tested animal. Sound levels that accounted for average sensation level as well as sound levels that accounted for age-related cochlear synaptic degeneration were used. As a whole, the results of this study should aid in unveiling the relationship of neural AM processing and behavioral AM perception in aging.

## 2. Methods

### 2.1. Animals

Twelve young (3-11 months; mean b.w.: male = 264 g and female = 183 g) and 14 aged (20-24 months; mean b.w.: male = 408 g and female = 242 g) Fischer-344 (F344) rats obtained from Taconic (NIA colony) were used. All animals were housed in the animal care facility during the period of this study in a relatively quiet and standard condition. They were also maintained on 12-hour light and 12-hour dark cycle (light on at 6:00 and off at 18:00) with water and food ad libitum. Behavioral experiments were performed during the light phase of the light-dark cycle, mainly in between 13:00 and 18:00. All protocols were approved by the Purdue Animal Care and Use Committee (PACUC-1111000167).

### 2.2. Behavioral tests (ASR and PPI)

#### 2.2.1 Setup and experimental procedure

All behavioral tests were performed in a sound attenuating cubicle (Med Associates) within a larger anechoic chamber (Industrial Acoustics). During the testing procedure, animals were placed on a grid rod animal holder on a motion-sensitive platform. Animals’ startle responses were detected and transduced via an amplifier connecting to a TDT RZ6 system and the com-puter. The vertical movement of the platform, which resulted from a startle reaction, was converted into a voltage signal by a transducer.

Startle responses were measured from the beginning of each trial to 1.5 s after the offset of the SES. Acoustic stimuli, including background sounds and prepulses, were generated by a TDT RZ6 system and presented via a Fostex (FT28D Dome Tweeter) speaker. The SES was also generated by the same TDT system and presented through a high frequency neodymium compression driver (BMS speaker). Both speakers were placed behind the animal holder. Stimulus presentation and response acquisition were manipulated by custom-written scripts using RPvdEx and MATLAB (MathWorks). Calibration of the apparatus was carried out for frequencies 1-20 kHz using a 1/2” Bruel & Kjaer microphone connecting to Nexus preamplifier and an os-cilloscope (Tektronix). The microphone was placed inside the animal holder at the middle of the cage, as recommended by the manual of Med Associates, during the process of sound calibration.

For every animal that has not performed any behavioral PPI test before, each of them was habituated to stay in the animal holder for 5-10 min for 3 successive days [68]. After 3 days of habituation, animals were then proceed to perform an 8 kHz pure tone detection task or AMF discrimination task. Each animal completed only one task (about 60 min) on one test day. A complete task encompassed a total of 3 phases, which were named as phase 0, 1 and 2. In summary, phase 0 is an acclimation period for animals to adapt to the animal holder, phase 1 is for habituation and association, and phase 2 is the period in which the detection or discrimination task used for analysis was carried out.

#### 2.2.2 8 kHz pure tone detection task

Animals’ abilities in detecting 8 kHz pure tones in a quiet background were tested using prepulses of 8 kHz pure tones at sound levels of 25-75 dB SPL in 10-dB difference. In phase 0, animals underwent acclimation for 5 min. In phase 1, 30 trials of SES alone were performed for animals to habituate to around 60 % of their initial startle responses [68]. Wideband noise of 20 ms duration with zero rise fall times was used as the SES. The intensity of the SES was set at 105 dB SPL for young animals and 115 dB SPL for aged animals. The interval between the onset of each trial was randomized between 15 and 30 sec so that animals could not estimate the appearance of a SES. Phase 2 contains trials with a SES alone (served as positive controls), trials with a prepulse placed before a SES and trials with a prepulse alone (served as negative controls). The prepulses were 8 kHz pure tones with a duration of 50 ms (5 ms rise fall times). The intensity of a prepulse in each trial was pseudorandomized between 25 and 75 dB SPL (10-dB gap). As each type of prepulse intensity repeated 9 times within one complete task, a total of 72 trials were consisted in phase 2. Similar to phase 1, the intertrial interval in phase 2 was also randomized between 15 to 30 s.

Behavioral 8 kHz detection threshold was estimated for each animal by comparing the ASR or RMS ratio measurements of no prepulse to the ASR or RMS ratio measurements of 8 kHz prepulses at various sound levels. Significant decreases in the ASR or RMS ratio measurements of prepulses from those of no prepulse were quantified using a one-sided t-test [41]. The minimum sound levels that elicited a significant decrease in both of the measurement were averaged. This mean threshold was then taken as the behavioral 8 kHz detection threshold for the particular animal.

#### 2.2.3 AMF discrimination task

AMF discrimination task was performed in a background of SAM tones. An 8 kHz carrier (200 ms) with 128 Hz AMF at 100, 50 or 25 % AM depth was presented as a background tone throughout the task. This SAM tone was repeated multiple times (about 12-27 times) before a prepulse and a SES were presented (Fig. 1). In phase 0, the background SAM tone was presented at 1 /s for 5 min to allow animals to acclimate to the animal holder and the background sounds. Phase 1, consisted of 20 trials, was used to habituate animals in associating the prepulse, which has an AMF different from the background, with a SES. In these 20 trials, the AMF of the prepulse was set at the highest or lowest AMF (depending on the range of the AMF that was tested in Phase 2) and presented alternatively. Fifty milliseconds after the prepulse (200 ms) offset, the SES was released. The intertrial interval was randomized between 15 and 30 s. The background AM tone was played during the 15-30 s interval but became silent for 2.6 s after the generation of a SES. The background AM tone was then resumed at the start of the next trial. Phase 2 contained a total 81 trials (each trial type repeated 9 times) and was used to measured PPI for AMF discrimination. The AMF of the prepulse was varied from trial to trial to test animals’ abilities in discriminating it from the background AMF. The startle magnitude was expected to be smaller if animals could discriminate the prepulse’s AMF from the background. In contrast, if animals could not discriminate the prepulse’s AMF from the background, the loud noise should trigger a relatively larger startle response. All the trials in phase 2 could be categorized into four conditions: (1) background only (negative control); (2) background and prepulse (negative control); (3) background and SES (positive control); and (4) background, prepulse and SES. Conditions (1) and (2) were negative controls because no startle response should be induced in these two conditions. Condition (3) served as a positive control since it contained a SES with no prepulse and a large startle response should be triggered. In condition (4), reduced startle response was expected if animals were able to discriminate a change in AMF from the background. The AMFs that were tested in both young and aged animals includes 16, 32, 64, 256, 512, 1024 Hz (*±*3-to *±*1-octave away from 128 Hz). A narrower AMF range was also tested in young animals and the AMFs are 45, 64, 90, 181, 256 and 362 Hz (*±*1.5-to *±*0.5-octave away from 128 Hz). The background SAM tones was randomly presented between 12 to 27 times (at 1/s for 12-27 s) from trial to trial in order to remove any other possible cues that could be used by animals to predict the SES. The only cue that should be used by animals to predict the SES would be based on their abilities to distinguish a change in AMF from the background’s AM. Each animal repeated the same PPI behavioral test for 2 times to confirm consistency. Overall, the experimental procedure, stimulus presentation and parameters for AMF discrimination task were designed by referring to the published literature [68, 56, 15].

**Figure 1:**
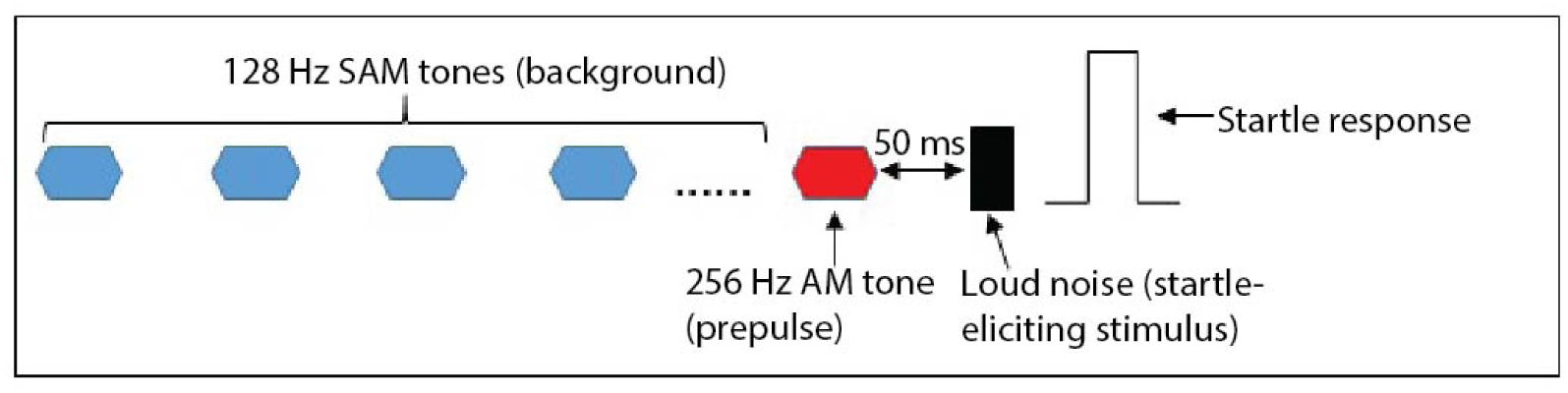
Presentation of background sounds, prepulse and startle-eliciting stimulus in a typical trial of the PPI behavioral task for AMF discrimination. The schematic shows an example of a PPI trial with multiple 128 Hz SAM tones presented in the background and a 256 Hz SAM tone used as a prepulse placing right before a startle-eliciting stimulus.

In term of stimulus intensity, the background and the prepulse levels were set at 85 dB SPL for aged animals and 75 dB SPL for young animals. This 10-dB difference in the sound level used in young and aged animals accounted for the average difference in sensation level at 8 kHz for young and aged animals [49]. In addition, for the first set of AMFs at 100 or 50 % AM depth, we also tested young animals using sound levels that matched to the aged’s median ABR tone 8 kHz wave I amplitude at 85 dB SPL in order to attain equivalent peripheral neural activation. This accounted for cochlear synaptopathy and/or neuropathy as well as age-related differences in hearing thresholds [60]. In this case, the average sound intensity was approximately 57.2 +/- 5.1 dB SPL in the young based on the measurement of tone 8 kHz ABR wave I amplitudes, which would be about 30 dB sensation level.

#### 2.2.4. Startle response measurements and PPI calculation

Animal startle responses were recorded by the platform and then filtered off-line with high-pass at 2 Hz and low-pass at 50 Hz. After filtering, a typical startle response has a specific waveform as shown in Figure 2. Two different methods were used to measure ASR responses [23]: (1) ASR magnitude: the maximal peak-to-peak amplitude of transient voltage occurring within 300 ms after the offset of the SES; (2) ASR root mean square (RMS) ratio: the RMS of the startle response (t_ASR_, corresponding to a -100 to +200 ms window relative to the first peak that occurred within 300 ms after the offset of the SES) over the RMS of the baseline (t_NF_, ref. Fig. 2). The measured mean ASR amplitude or mean RMS ratio for each trial type was estimated as the average of all the ASR amplitudes or the RMS ratios after the highest and lowest values were excluded [67]. This is to remove any possible outliers as well as reduce variability of the responses. The percent of PPI (i.e. the percent of startle magnitude reduced by the prepulse as compared to the positive control) for each trial type was calculated using the below formula:

PPI % = [1-(ASR magnitude or RMS ratio to prepulse - baseline)/ (ASR magnitude or RMS ratio of startle only-baseline)] x 100 %.

**Figure 2:**
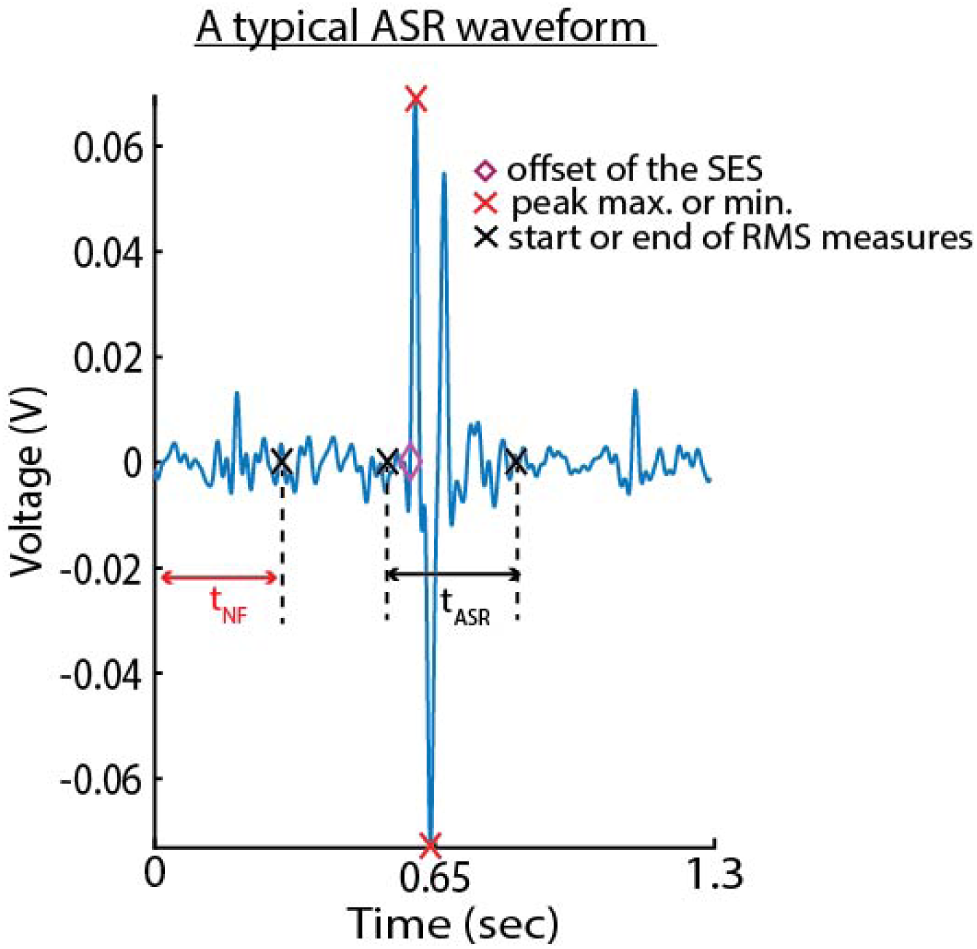
A typical acoustic startle response (ASR) waveform with distinct peaks and troughs that are above or below the noise floor (NF). The schematic shows an example of an ASR waveform obtained from a PPI trial. The offset of the startle-eliciting stimulus (SES), the start and end for root-mean-square (RMS) measures are labeled on the plot. For RMS ratio calculation, the time window of an ASR response is denoted by t_ASR_ while t_NF_ indicates the time window used for the noise floor. Both t_ASR_ and t_NF_ are 300 ms in duration.

Magnitude or RMS ratio of baseline was measured from negative controls (trials with no SES) while startle only was measured from positive controls (trials of background and loud noise with no prepulse). A PPI % value that is close to or at 0 indicates that the prepulse does not have an inhibitory effect on animals’ startle responses, which also indicates that animals could not discriminate the prepulse from the background. However, a PPI % that is near to 100 % indicates an almost complete inhibition of startle responses by the prepulse.

### 2.3. Auditory evoked potentials

The experimental protocols used for ABR and EFR recordings were similar to previously described details in Parthasarathy and Bartlett (2012). All recordings were performed in a 9’x9’ double-walled anechoic chamber (Industrial Acoustic Corporation). The animals were anesthetized using isofluorane at 4 % and later maintained under 1.5-2 % isofluorane for placing the electrodes. Subdermal needle electrodes (Ambu) were placed on the animals’ scalps in a two-channel configuration. For channel 1, a positive electrode was placed along the midline of the forehead in the the Cz to Fz position. For channel 2, another positive electrode was placed horizontally along the interaural line, which is above the location of the inferior IC. The negative electrode was placed under the ipsilateral ear, along the mastoid, while the ground electrode was placed in the nape of the neck. Electrode impedance was confirmed to be less than 1 kΩ by testing with a low-impedance amplifier (RA4LI, Tucker Davis Technologies or TDT). Before taking off isofluorane, the animals were injected (intramuscular) with dexmedetomidine (Dexdomitor, 0.2 mg/kg), an α-adrenergic agonist acting as a sedative and an analgesic. Recording was then started after a 15-min waiting time for the effects of isofluorane to wear off. The animals were maintained in an unanesthetized and immobile condition during the whole session of recording.

Tone 8 kHz ABRs were recorded using brief 8 kHz pure tones of 2 ms duration (0.5 ms cos^2^ rise/fall time), alternating polarity and presenting at 26.6/sec. The acquisition window was set to 30 ms and each ABR was acquired as an average of 1500 repetitions (750 each polarity). Stimulus intensity of the pure tone was decreased from 95 dB SPL to 15 dB SPL in 5-dB steps. This enabled us to obtain the animal’s hearing threshold at 8 kHz as well as the magnitude of wave I at each sound level, which was used as an indicator for the amount of activated ANFs. The median of tone 8 kHz ABR wave I amplitudes at 85 dB SPL from aged animals was used for stimulus intensity matching of peripheral activation in young animals. Sinusoidally amplitude modulated (SAM) tones with a 8 kHz carrier were used as acoustic stimuli for EFRs. At 100 %, 50 % or 25 % modulation depth, the AMF of the SAM tones was systematically increased from 16 to 2048 Hz in 0.5-octave steps to generate the tMTF. The stimulus intensity was set at 75 dB SPL for young animals and 85 dB SPL for aged animals. In young animals, sound levels that matched to the aged’s median ABR tone 8 kHz wave I amplitude at 85 dB SPL were also recorded.

All stimuli were presented free-field to the right ear of the animal at a distance of 115 cm from a speaker (Bower and Wilkins DM601). Stimuli were generated using SigGenRP (TDT) at a 100-kHz sampling rate. Stimuli presentation and response acquisition were conducted using BioSig software (TDT). Waveforms were converted to sounds and delivered through a multichannel processor (RX6, TDT) via the speaker. Digitized response waveform was recorded with a multichannel recording and stimulation system (Rz5, TDT). Responses were analyzed with BioSig and a custom-written program in MATLAB.

All collected EFRs were low-pass filtered at 3000 Hz. EFRs were also high-pass filtered at 12 Hz for AMFs of 12-24 Hz, 30 Hz for AMFs of 32-64 Hz and 80 Hz for AMFs faster than 90 Hz. Filtered data were then exported as text files and analyzed using custom-written MATLAB scripts. Fast Fourier transform (FFT) were performed on time-domain waveforms from 10 to 190 ms relative to stimulus onset to exclude transient auditory brainstem responses at the beginning. The maximum magnitude of the evoked response at one of the three frequency bins (3 Hz/ bin) around AMF was measured as the peak FFT amplitude. The noise floor was calculated as the average magnitude of five frequency bins above and below the central three bins. A peak response was taken to be significantly above noise level if the FFT amplitude was at least 6 dB above the noise floor for the slower AMFs and at least 10 dB above the noise floor for AMFs faster than 64 Hz to account for the sharply decreasing noise floor.

### 2.4. Statistical analysis

Repeated measures ANOVAs (rmANOVAs) were performed to compare ASR responses or FFT amplitudes of young and aged groups as well as across different stimulus conditions using custom written scripts in SAS (Proc MIXED, SAS Institute, Cary, NC, USA). Main effects and interactions effects of each factor were analyzed based on comparisons of least squares (LS) means. Data distributions were checked for normality using normal probability plots of the residuals (proc UNIVARIATE). The differences in LS means with a confidence level of 95 % was used when reporting significant differences. LS means +/- standard error of mean (SEM) are shown in the figures.

## 3. Results

### 3.1. 8 kHz tone detection in a quiet background

Prepulses of 8 kHz pure tones at sound intensities of 25-75 dB SPL, in 10-dB difference, were used to test animals’ hearing sensitivities at 8 kHz. The growth of PPI as a function of sound level, i.e. PPI values increased as 8 kHz prepulse intensity increased, was observed in young and aged animals as shown in Fig. 3. For almost all of the sound levels, young animals had larger PPI values than old animals although age-related differences were not statistically significant. For each age group, PPI values at higher sound levels were significantly larger than PPI values at lower sound levels, e.g. 75 *>* 35 db SPL. Table 1 shows sound levels with PPI that are significantly different from each other in young and aged animals for each of the measurement. In addition, SEM of aged animals tended to be larger at lower sound levels (25-45 dB SPL). This indicates that young animals were more behaviorally consistent at perceiving 8 kHz tones at lower sound levels because of having better hearing sensitivity. In young animals, the mean PPI values at each sound level were significantly larger than 0 when tested using a t-test. However, the mean PPI values were significantly larger than 0 in aged animals at higher sound levels. Statistical analysis using rmANOVA revealed a significant main effect of sound level for the measurement of ASR magnitude (F = 17.52, p *<* 0.05) and ASR RMS ratio (F = 13.05, p *<* 0.05). However, no significant age or age*sound level effect was observed for both measurements.

**Figure 3:**
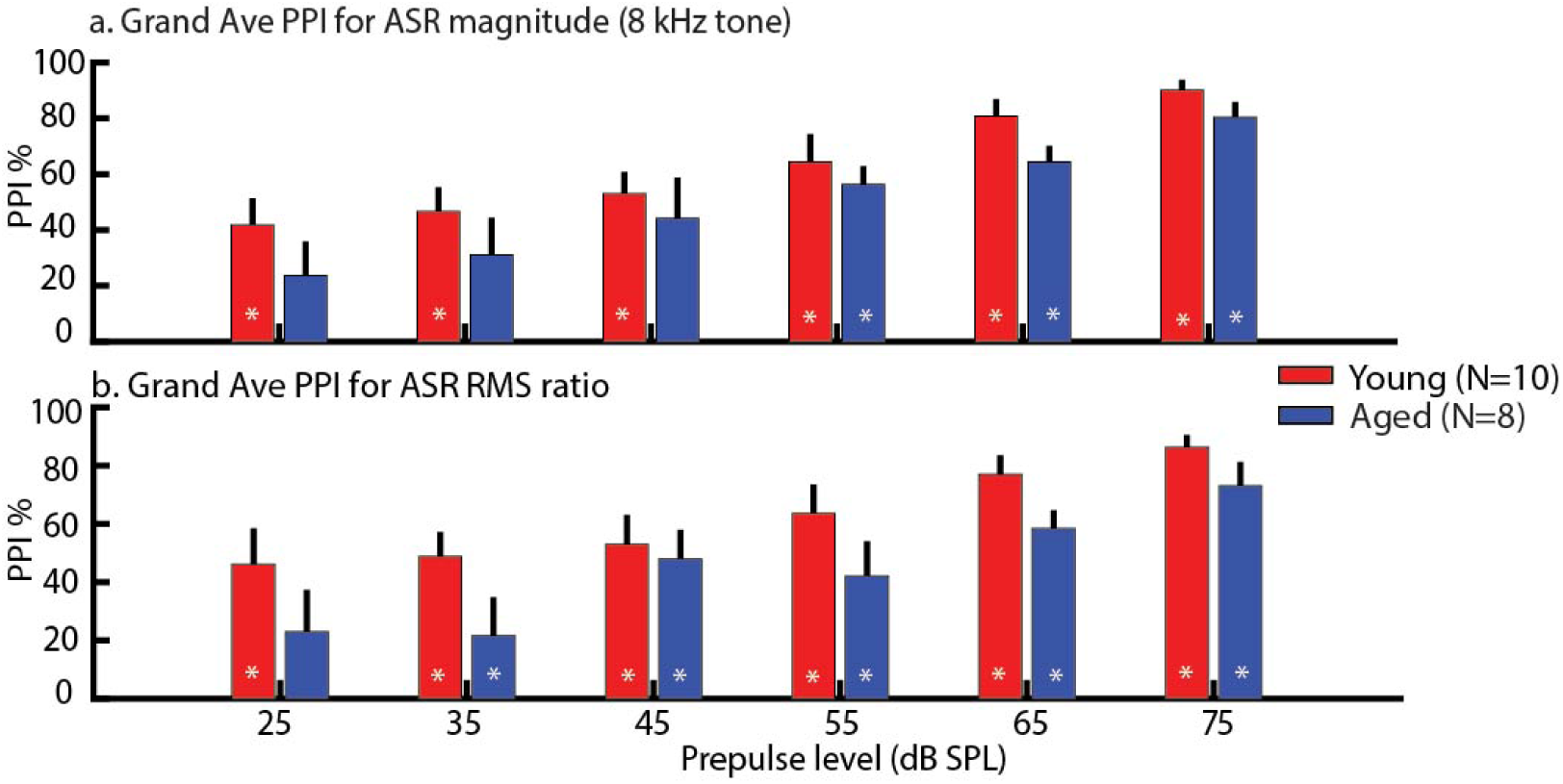
Prepulse inhibition (PPI) using prepulses of 25-75 dB SPL 8 kHz pure tones in a quiet background showed similar growth in PPI as sound intensity increased in young and aged animals. PPI values of higher sound intensities were larger than those of lower sound intensities. The black asterisks indicate p *<* 0.05 for PPI comparison between age groups and at the same sound level. The white asterisks in bars indicate p *<* 0.05 for mean PPI not equal to zero using a t-test. All statistically significant differences were obtained using least squares means comparison from rmANOVA and PPI comparison between sound levels within an age group is summarized in Table 1.

**Table 1:**
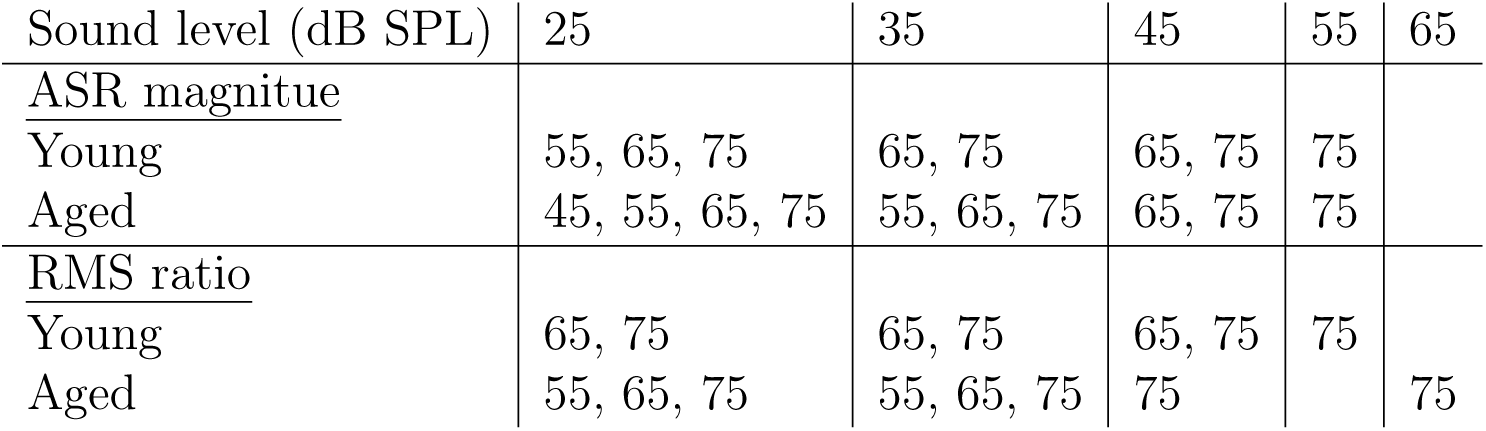
For 8 kHz prepulse detection, PPI values of lower sound levels were mostly significantly different from PPI values of higher sound levels. This table shows sound levels with PPI that are significantly different from each other within each age group according to the results of rmANOVAs for Figure 3.

| Sound level (dB SPL) | 25 | 35 | 45 | 55 | 65 |
| --- | --- | --- | --- | --- | --- |
| <u>ASR magnitue</u> |  |  |  |  |  |
| Young | 55, 65, 75 | 65, 75 | 65, 75 | 75 |  |
| Aged | 45, 55, 65, 75 | 55, 65, 75 | 65, 75 | 75 |  |
| <u>RMS ratio</u> |  |  |  |  |  |
| Young | 65, 75 | 65, 75 | 65, 75 | 75 |  |
| Aged | 55, 65, 75 | 55, 65, 75 | 75 |  | 75 |

Behavioral 8 kHz detection threshold estimation using the measurements of ASR and RMS ratio was performed for each animal. Young animals generally have lower 8 kHz detection thresholds than aged animals. The mean 8 kHz detection threshold of the young was 39.5 +/- 0.2 dB SPL while the mean 8 kHz detection threshold of the aged was 61.9 +/- 0.17 dB SPL. How-ever, these thresholds were higher than the 8 kHz hearing thresholds obtained from ABRs elicited by brief 8 kHz tones. The measured mean tone 8 kHz ABR threshold for the young was 25.5 +/- 0.04 dB SPL and for the aged was 37.2+/- 0.09 dB SPL. Statistical comparisons of hearing thresholds for age vs. young or behavior vs. ABR were performed using rmANOVAs. The results show main effect of Age (F = 12.44, p *<* 0.05) and Measure type (F = 22.61, p *<* 0.05) but no significant interaction effect.

### 3.2. Behavioral discrimination of AMFs

### 3.3. In young animals

The first set of frequencies tested in young animals for AMF discrimination includes the range of 16-1024 Hz with 1-octave difference. Each AMF is 1, 2 or 3 octaves higher or lower than 128 Hz AM. The same AMF discrimination task was performed by fixing AM depths of all SAM tones at either 100, 50 or 25 %. The PPI results obtained with these three AM depths using either ASR magnitude or RMS ratio were shown in Figure 4. When comparing PPI values among different AM depths but at one single AMF, higher inhibition was observed for larger AM depths compared to smaller AM depths, e.g. 100 % *>* 50 % *>* 25 %. Statistical significance for PPI values being higher at larger AM depths compared to smaller AM depths was observed at most AMFs. In addition, when comparing PPI values across different AMFs but within the same AM depth, a trend of higher PPI was observed at AMFs that were further away from 128 Hz for 50 and 25 % AM depths. At 25 % AM depth, grand average PPIs of almost all the tested AMFs generally had larger SEMs. This indicates that behavioral variability among young animals in AMF discrimination increased when AM depth reduced. According to the results of t-tests, the mean PPI values at each AMF at 100 and 50 % depth were all significantly different from 0 indicating significant inhibitory effects. In contrast, the mean PPI values at 25 % depth were not significantly different from 0 at most AMFs except 1024 Hz. In addition, a significant main effect of AM depth was obtained from rmANOVA for the measurements of ASR magnitude (F = 10.51, p *<* 0.05) and RMS ratio (F = 14.54, p *<* 0.05).

**Figure 4:**
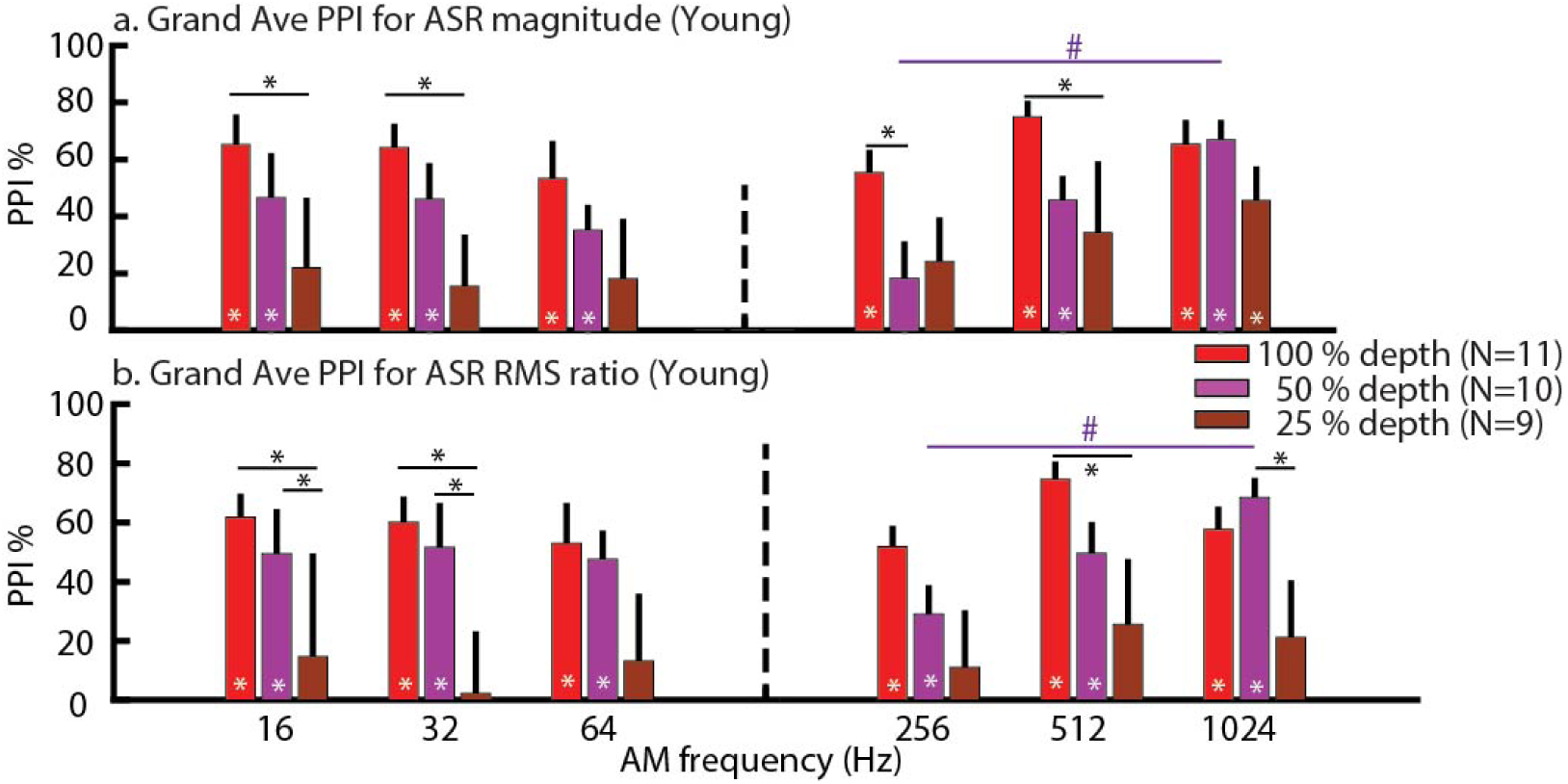
In young animals, PPI values were higher for larger AM depths compared to lower AM depths (e.g. 100 % *>* 50 % *>* 25 %) at various AMFs (16-1024 Hz in 1-octave difference). For 50 % AM depths, PPI tended to increase as AMFs were further away from 128 Hz. The black asterisks indicate p *<* 0.05 for PPI comparison between different AM depths within the same AMF while the pound signs indicate p *<* 0.05 for PPI comparison between different AMFs but within the same AM depth. All statistically significant differences were obtained using least squares means comparison from rmANOVA. The white asterisks in bars indicate p *<* 0.05 for mean PPI not equal to zero using a t-test.

The second set of frequencies tested on the young includes the range of 45-362 Hz separated in 0.5-octave difference. Each AMF is 0.5-, 1- or 1.5-octave away from 128 Hz AM. In Figure 5, PPI values at 100 % depth were relatively higher than 50 % depth. When comparing PPI across different AMFs at 50 % AM depth, a trend of increased PPI was observed when AMFs were further away from 128 Hz. Moreover, for 50 % AM depth, grand average PPI of most AMFs had larger SEM indicating variability among young animals in AMF discrimination increased as AM depth reduced. The mean PPI values were significantly larger than 0 for almost all AMFs at 100 % depth but not for 50 % depth. According to rmANOVA, there is a significant main effect of AM depth for both ASR magnitude measurement (F = 17.69, p *<* 0.05) and RMS ratio measurement (F = 11.74, p *<* 0.05).

**Figure 5:**
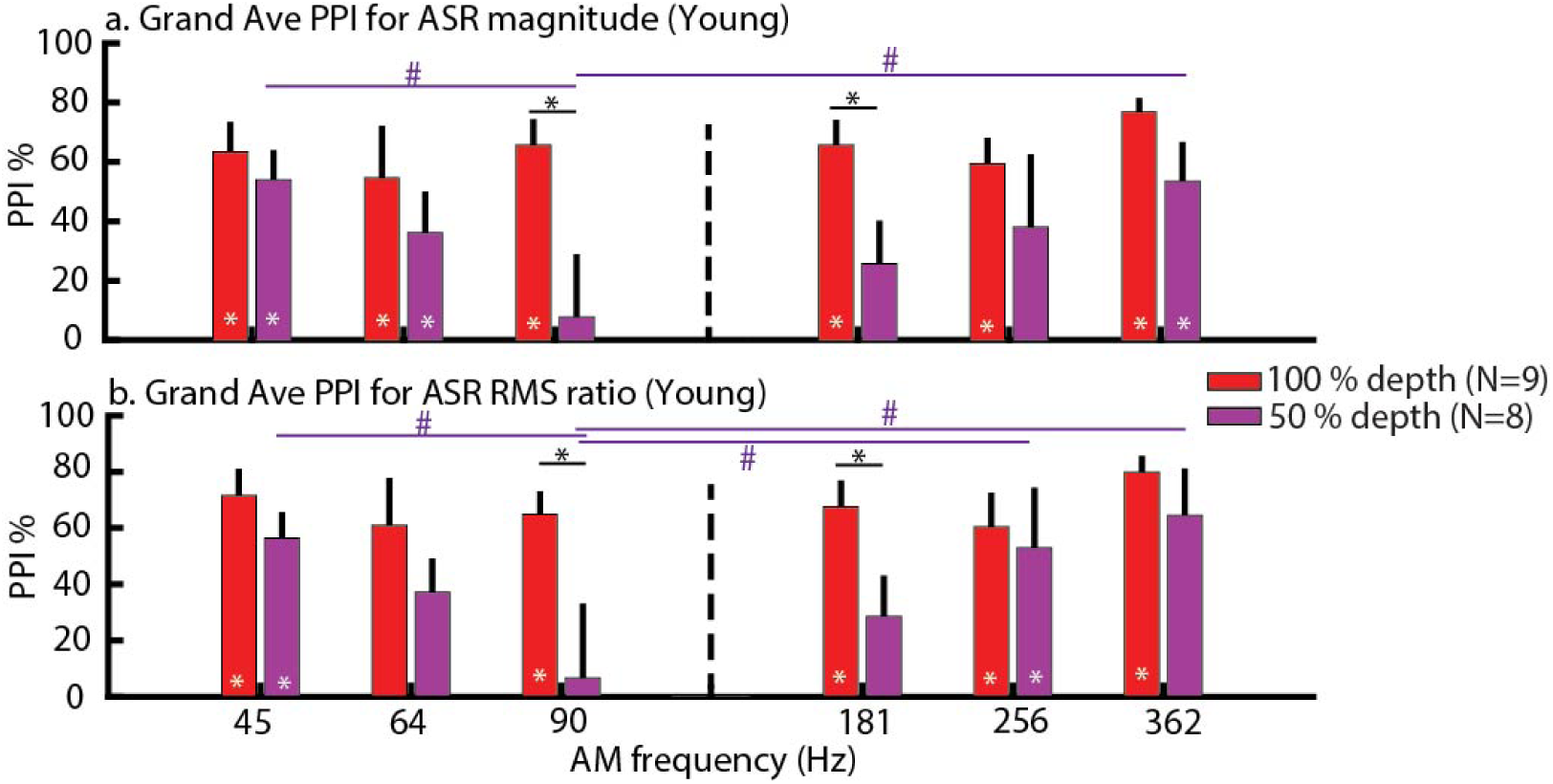
A trend of higher PPI was observed for 100 % AM depth compared to 50 %. The PPI results were obtained from a more difficult task in which the AMF range was set at 0.5-1.5 octave away from 128 Hz. Indications for the asterisk and the pound signs are similar to Figure 4.

### 3.4. Young vs. aged animals

AMF discrimination was tested in young and aged animals using stimulus intensity of either 75 (young) or 85 db SPL (aged). The tests were performed at either 100 or 50 % AM depth. Young animals were also tested at sound levels (an average of about 55.3 db SPL) that matched to the aged median tone 8 kHz ABR wave I amplitude to achieve equivalent peripheral neural activation. This accounted for cochlear synaptopathy and/or neuropathy [60] as well as age-related differences in hearing thresholds because ABR wave I amplitude reflects the amount of activated and synchronized auditory neurons [55, 9]. Figure 6 shows the results of PPI obtained at 100 % AM depth. There was a trend of aged PPI values at 85 dB SPL being lower than PPI of the young at 75 dB SPL and at matched peripheral activation. Young PPI values at 75 dB SPL and at matched peripheral activation were similar except at 1024 Hz AMF. Statistical analysis using rmANOVA revealed significant main effect of AMF for PPI measured with ASR magnitude (F = 4.1, p *<* 0.05) and RMS ratio (F = 3.42, p *<* 0.05).

**Figure 6:**
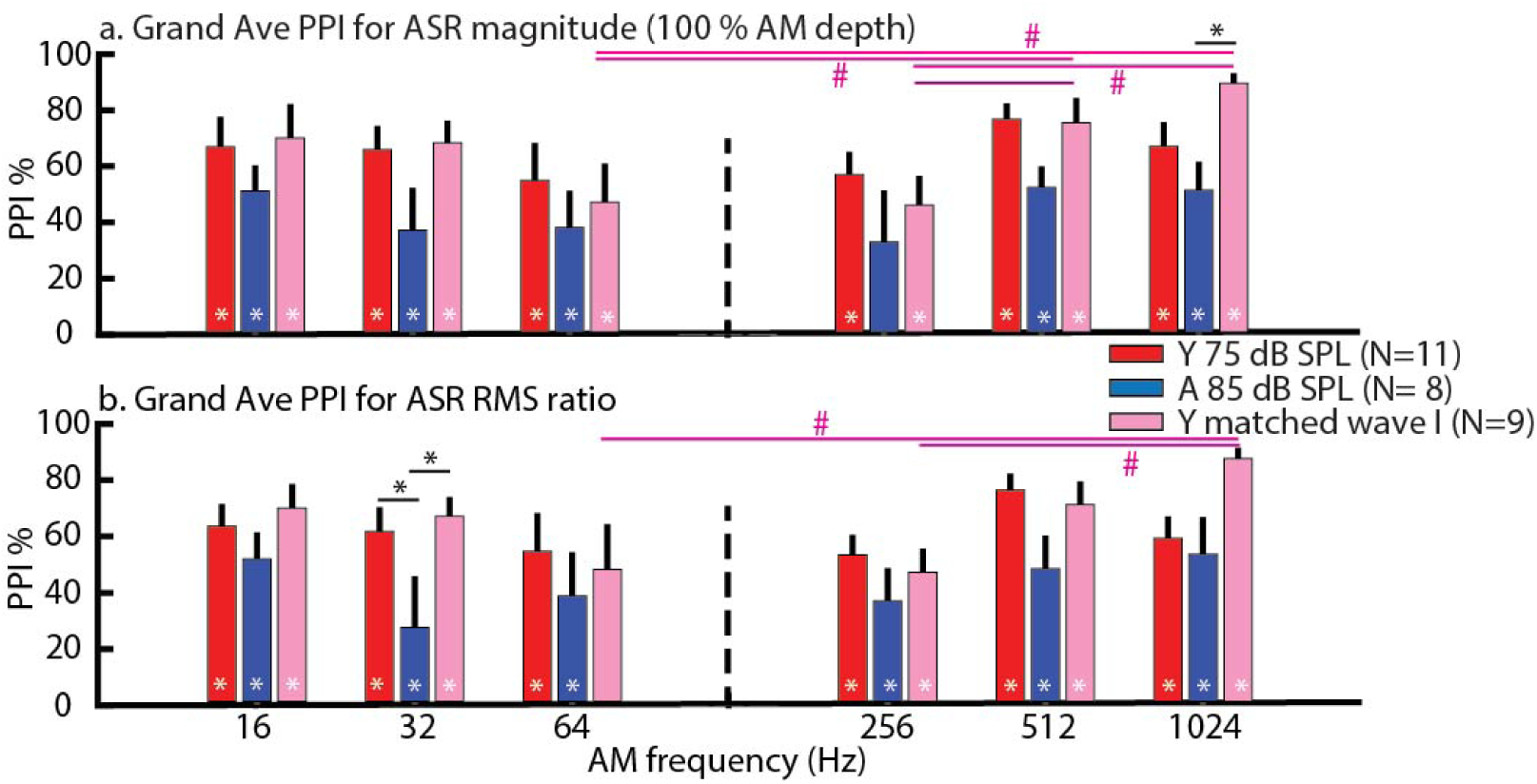
PPI was detectable in aged animals for almost all AMF differences for one octave spacing and 100% AM depth. There was a trend of aged PPI values being lower than the young at 75 dB SPL and at matched peripheral activation. The pound signs indicate p *<* 0.05 for PPI comparison between different AMFs within the same age group. All statistically significant differences were obtained using least squares means comparison from rmANOVA. In the legend, Y indicates young animals while A indicates aged animals. The white asterisks in bars indicate p *<* 0.05 in t-test for mean PPI not equal to zero. In the legend, Y indicates young animals while A indicates aged animals.

Figure 7 shows the results of PPI obtained at 50 % AM depth. In the young 75 dB SPL, PPI values were generally smaller than for 100 % depth (cf. Fig 6), but still showed PPI significantly higher than zero. By contrast, the PPI responses for the aged 85 dB SPL and the young with peripheral matching were not significantly above zero at some AMFs (e.g. 16, 256 and 512 Hz). When AM depth reduced to 50 %, AMF discrimination abilities for the aged at 85 dB SPL and the young at matched peripheral activation reduced, especially at 256 Hz AMF. According to rmANOVAs, there was a significant main effect of AMF obtained from rmANOVAs for PPI measured using the ASR magnitude method (F = 6.71, p *<* 0.05) and the ASR RMS ratio method (F = 7.55, p *<* 0.05). The rmANOVA results for the ASR RMS ratio also showed a significant main effect of Age (F = 9.28, p *<* 0.05).

**Figure 7:**
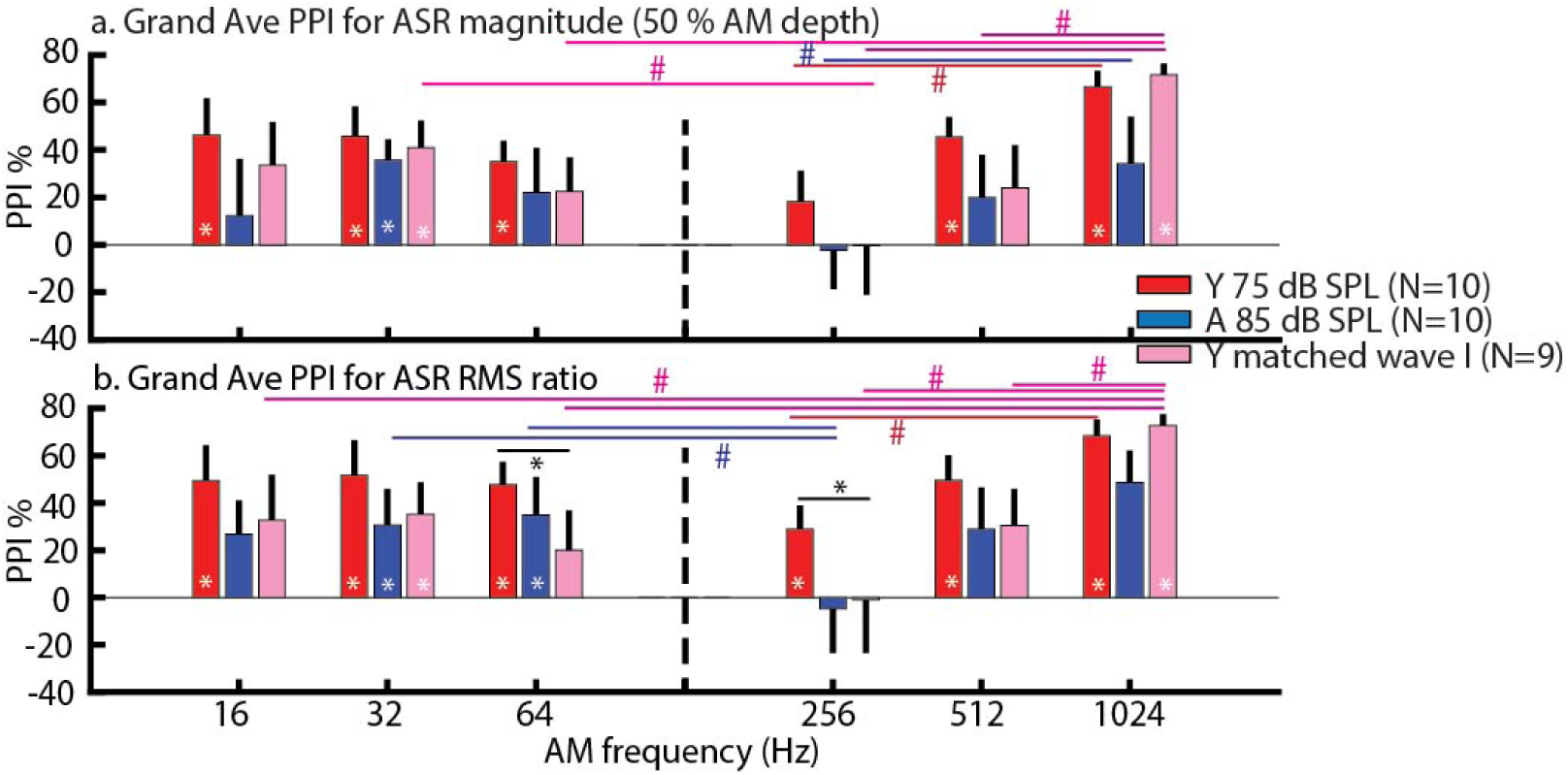
For AMF discrimination at 50 % AM depth, a trend of higher PPI values in young animals (75 dB SPL) across AMFs was observed. Aged animals had PPI close to baseline or in negative values especially when responses were measured using RMS ratio. PPI values of young animals at 75 dB SPL or matched wave I were mostly not significantly different from the aged. The black asterisks indicate p *<* 0.05 for PPI comparison between age groups but at the same AMF. All statistically significant differences were obtained using least squares means comparison from rmANOVA. The white asterisks in bars indicate p *<* 0.05 in t-test for mean PPI not equal to zero. In the legend, Y indicates young animals while A indicates aged animals.

### 3.5. Electrophyiological responses for AMF perception

Electrophyiological responses elicited by AMFs ranging from 16-2048 Hz were recorded in both young and aged animals via EFRs using 8 kHz tone carriers (Fig. 8a). Sound levels were set at 75 dB SPL for the young and 85 dB SPL for the aged, which has been shown to evoke peak EFR responses in most animals [47]. Fig. 8a shows EFRs of tMTFs with 100, 50 or 25 % AM depth in young and aged animals. At 100 % AM depth, the young EFRs were generally higher than the aged even though the stimulus level used in the aged was 10 dB SPL louder. For aged animals, their EFRs at 100 % AM depth were similar to the young EFRs at 50 % AM depth. Moreover, the aged EFRs at 50 % AM depth were also similar to the young EFRs at 25 % AM depth. However, when EFRs of tMTFs were recorded at equivalent peripheral activation, the aged EFRs at 100 % AM depth were significantly higher than the young EFRs at 100 % AM depth (Fig. 8b). Although differences were smaller, the aged EFRs at 50 % AM depth were still significantly larger than the young EFRs at 50 % AM depth. According to statistical analysis using rmANOVA for EFRs recorded at equivalent peripheral activation, the main effects of age and AMF as well as their interaction effect were statistically significant (p *<* 0.05). At 100 % AM depth, the F-values of age and AMF main effects were 19.97 and 52.92, respectively. The interaction effect of age*AMF had an F-value of 5.68. For 50 % AM depth, the F-values of age and AMF main effects were 6.68 and 179.12, respectively while the F-value for the interaction effect of age*AMF was 2.13. We did not perform statistical analysis for EFRs in Fig. 8a because the emphasis was to observe the trends and how EFRs of tMTFs with different AM depths were distinct or overlapped.

**Figure 8:**
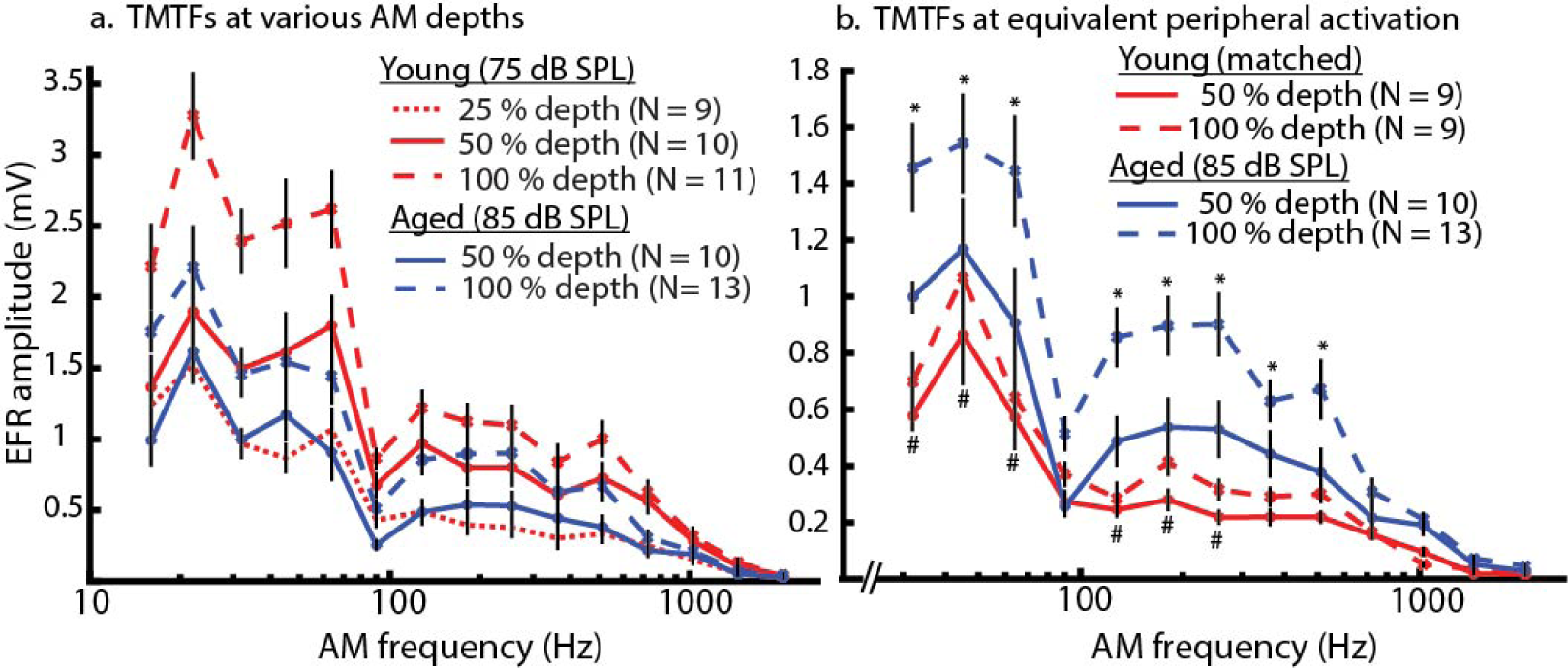
Young animals’ EFR amplitudes were generally larger at 75 dB SPL compared to aged animals at 85 dB SPL but their EFR amplitudes were lower than the aged at equivalent peripheral activation. (a) EFRs of temporal modulation transfer functions (tMTFs) with 100, 50 or 25 % AM depth recorded in young and aged animals, respectively. (b) EFRs of tMTFs with 100 or 50 % AM depth recorded in both age groups at matched peripheral activation. The asterisks indicate p *<* 0.05 for comparison of EFR amplitudes between young and aged animals for tMTFs with 100 % AM depth while the pound signs indicate p *<* 0.05 for comparison of EFR amplitudes between young and aged animals for tMTFs with 50 % AM depth. All statistically significant differences were obtained using least squares means comparison from rmANOVA.

### 3.6. Relationship of EFRs and behavioral PPI

To identify the relationship between neurophysiological responses and behavioral AMF discrimination at each of the tested AMFs, changes in each of these measures due to a change in temporal salience of AM depth were compared simultaneously. The changes in behavioral PPI or the changes in EFR amplitudes as temporal salience of AM depth dropped from 100 to 50 % were measured at each of the tested AMF and in each age group. As shown in Figure 9, changes in PPI values were plotted on the left ordinate while changes in EFR amplitudes were plotted on the right ordinate. The changes in PPI values (ΔPPI) were measured as PPI % at 100 % AM depth minus PPI % at 50 % AM depth from the same animals. The changes in EFRs (EFR ratio) were measured as EFR amplitudes at 50 % depth divided EFR amplitudes at 100 % depth from the same animal as well.

**Figure 9:**
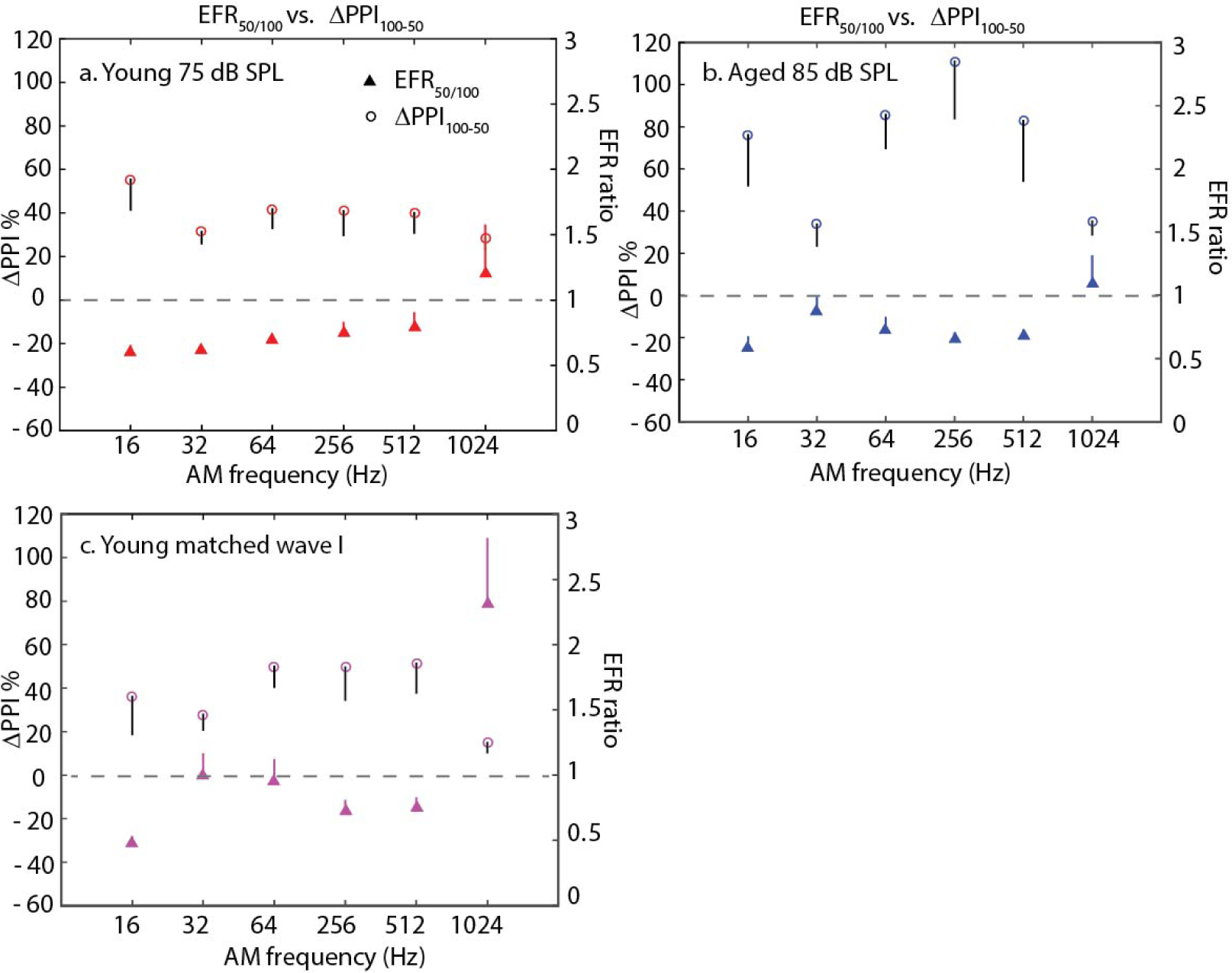
Greater changes of behavioral PPI values compared to changes of EFRs in aged animals when salience of AM depth reduced. Left ordinate indicates the measure of ΔPPI, which is the difference of PPI % at 100 % AM depth versus 50 % AM depth. Right ordinate indicates the measure of EFR ratio, which is the ratio of EFR amplitude at 50 % AM depth versus 100 % AM depth. The change in PPI value or EFR amplitude due to a change in AM depth was measured from the same animal in (a) young animals (75 dB SPL), (b) aged animals (85 dB SPL), and (c) young animals at equivalent peripheral activation. The paired changes were then averaged and the means of paired differences +/- SEM were plotted.

For young animals (75 dB SPL), consistent smaller changes in EFRs and PPIs due to a decrease in stimulus AM depth were observed. This indicates that their abilities in AMF discrimination and EFR responses to the tested AMFs were not much affected by a reduction in AM depth. For aged animal (85 dB SPL), the trend of EFR ratio over AMF behaved similarly to young animals (75 dB SPL) but their ΔPPIs were larger compared to young animals (75 dB SPL). There was a larger change in behavioral AMF discrimination performance due to a reduction in AM depth although changes in EFRs were relatively smaller. The trend observed in young animals seemed to hold even when they were tested at matched peripheral activation. The changes in behavioral PPI were slightly larger compared to those at 75 dB SPL. Overall, a smaller change in EFR correlated with a smaller change in behavioral PPI value in young animals at both 75 dB SPL and at equivalent peripheral activation. However, this correlation was no longer consistent in aged animals.

## 4. Discussion

### 4.1. Behavioral PPI audiometry versus ABRs

The paradigm of behavioral ASR and PPI has been used to assess auditory behavior in rodents [56, 65, 63, 42, 45, 19, 62, 41]. Using standard PPI techniques in the absence of a background sound, both younger and older animals exhibited PPI whose amplitude increased with increasing salience of the prepulse (Fig. 3). For a 25 dB prepulse, PPI was significantly larger than 0 in younger animals, comparable to their ABR thresholds and consistent with previous studies [42]. As expected based on the ABR thresholds, PPI magnitudes tended to be smaller in older animals for lower prepulse levels, but still grew with increasing level and achieved similar peak PPI. Therefore, animals of all ages tested exhibited the PPI behavior and to a similar degree.

### 4.2. Aging effects on PPI of ASR

Age-dependent reduction on startle responses elicited by acoustic stimuli in rodents, including F344 rats, have been reported in previous studies [56, 69, 45, 30, 38]. It has been suggested that age-related changes in ASR cannot be directly attributed to hearing loss because different ASR amplitudes were obtained from young adult rats of different strains with similar hearing sensitivities [56]. In our study, we observed comparable PPI values, especially at supra-threshold prepulse levels, for 8 kHz detection task in young and aged animals (Fig. 3). This is different that the reduction of PPI efficiency associated with aging reported in F344 rats by Rybaklo et al. (2012). At 100 % AM depth (Fig. 6), the aged and young had similar PPI values for AMF differences of 2-3 octaves. For 1 octave AMF difference, PPI tended to be reduced in the aged 85 dB SPL and the young with peripheral matching (Fig. 6). When AM depth salience decreased (Fig. 6), the observed age-related reductions of PPI further suggest a deficit in temporal processing leading to impaired perception.

### 4.3. AM frequency discrimination

Amplitude modulation is used by humans and animals to aid in auditory object formation [5, 3]. Many studies have used tMTFs as a measure of temporal acuity of the auditory system in psychoacoustic [71, 26, 1, 35] as well as in electrophysiological studies [12, 47, 52]. AM depth sensitivity as a function of AMF has been demonstrated as similar for rats [35] and other mammals, including humans [71] and chinchillas [27]. A progressive decrease in AM depth sensitivity (behavioral threshold became worse) of a noise carrier modulated between 5-2000 Hz were observed in rats [35] and rats having better AM depth sensitivity at AMFs of 10-60 Hz was also found to be similar to humans [71]. The behavioral tMTFs of humans [44], rats [35], barn owls [11] and chinchilla [57] showed a low-pass characteristic for AM detection resembling the electrophyiological tMTFs in F344 rats shown in this study (Fig. 8) and in our previous study [47]. For low modulation depths (25%), there was little evidence of discrimination in young animals for most AMFs. Despite this, PPI was evident for 1024 Hz AM (Fig. 4a), suggesting that AM discrimination even at low modulation depths (25%) is possible at AMF well above those that thalamic and cortical neurons can phase-lock to [34], suggesting that spectral cues and rate coding may be used. As task difficulty increased by reducing AM depth (Fig. 7), aged animals performed worse. Young animals tested at equivalent peripheral activation (55.3 dB SPL) performed better than the aged 85 dB SPL (Fig. 7) implying that peripheral activation by itself does not fully account for behavioral performance.

### 4.4. Correlation of behavioral auditory responses and the underlying neural responses

When the temporal salience of AM depth was decreased from 100 to 50% depth, the degree of the EFR phase-locking to the SAM stimuli decreased (Fig. 8 and 9). If EFR amplitudes have a strong link to behavioral performance, we expect that this should result in a decline in temporal perception (Fig. 9). When we compared changes in EFRs versus changes in behavioral PPI values due to a change in AM depth, Figure 9 reveals that both neuro-physiological and behavioral changes in young animals were correlated at 75 dB SPL as well as at softer sound levels (equivalent peripheral activation). A relative smaller change in behavioral PPI was associated with a relative smaller change in neural responses to SAM stimuli at the tested AMFs in the young 75 dB SPL. However, this correlation was no longer seemed to hold in the aged 85 dB SPL. A relatively smaller reduction in EFRs was observed to result in a larger decline in behavioral PPI in aged animals. This observa-tion is analogous to the findings of Xu and Gong (2014). When behavioral frequency difference limens (FDLs) and two-tone evoked frequency-following responses (FFRs) were measured in normal hearing young adults, they observed that frequency difference of two-tone, which was able to evoked FFRs, was smaller than behavioral FDL threshold [74]. Therefore, these and our results show that the neurophysiological measurements of EFRs or FFRs may be more sensitive than behavioral measurements because a smaller change in stimulus parameters can be detected physiologically but the response is not expressed behaviorally. Other behavioral tasks may be more sensitive, or it may be that phase-locking physiological measures are too sensitive [28]. These data also suggest that age-related degradation that exists beyond the auditory brainstem and midbrain could have a larger contribution to the decline in behavioral perception [73]. In addition, since we performed tone 8 kHz ABR wave I amplitude matching to achieve equivalent peripheral activation, which accounts for age-related increase of hearing threshold and age-related neuropathy/synaptopathy [60, 70], age-related decline in behavioral AMF discrimination should be due to more of a central effect and less to a peripheral effect.

In conclusion, we examined the relationship of behavioral AM perception and neurophysiological responses to similar stimuli by measuring PPI of ASRs and EFRs. The young behavioral performance in discriminating different AMFs dropped gradually as salience of AM depth reduced from 100 to 25 % depth. Comparable behavioral performances at AMFs 1-2 octaves away from 128 Hz were observed in young and aged animals when AMF spacing was larger and at 100 % AM depth. At 50 % AM depth, age-related decline of EFRs was smaller but aged animals’ AMF discrimination performance was highly compromised. When physiological and behavioral results were compared, the correlation of AM processing and AM perception were identified to be more consistent in the young, including even when peripheral activation was matched. Overall, the results reveal a larger age-related deficit in behavioral perception compared to auditory evoked potentials using similar SAM stimuli. This suggests that behavioral and physiological measurements should be combined to capture a more complete view on the auditory function and aid in identifying the localization of age-related auditory deficits.

